# Tobramycin enhances *Mycobacterium abscessus* fitness through *whiB7* induction

**DOI:** 10.1101/2025.10.01.679559

**Authors:** Jodi M. Corley, Jack H. Congel, Kelsey C. Haist, Alma E. Ochoa, Kenneth C. Malcolm, William J. Janssen, Jerry A. Nick, Katherine B. Hisert

## Abstract

Nontuberculous mycobacteria (NTM) are opportunistic pathogens that cause pulmonary disease (PD) in people with bronchiectasis and other chronic airways diseases. Difficulty treating and eradicating NTM-PD highlights the need for improved understanding of bacterial mechanisms to establish chronic infections. People with the genetic disorder cystic fibrosis (CF) develop bronchiectasis and are the population at highest risk of NTM-PD, caused mainly by *Mycobacterium avium* or *Mycobacterium abscessus* (*Mabsc*). The majority of people with CF (pwCF) and bronchiectasis develop chronic *Pseudomonas aeruginosa* airway infections. We hypothesized that antibiotics used to treat *P. aeruginosa* infections could enhance *Mabsc* persistence in the CF airway. Here we demonstrate that clinically relevant concentrations of tobramycin, which does not kill *Mabsc* but is frequently administered to pwCF with chronic *P. aeruginosa* infections, induced *Mabsc* expression of *whiB7*, a transcription factor that activates genes associated with resistance to host defenses. Tobramycin promoted *Mabsc* resistance to killing by hydrogen peroxide (H_2_O_2_) and survival in both human macrophages and mice. Deletion of *whiB7* increased *Mabsc* susceptibility to killing by H_2_O_2_, decreased ability of *Mabsc* to persist in macrophages, and disrupted ability of tobramycin to enhance *Mabsc* survival. Transcriptomic data defining the tobramycin associated WhiB7 regulon revealed differential gene expression of factors that could enhance *Mabsc* resistance to stress conditions such as those in found in the CF lung. Overall, our data indicate that administration of tobramycin to pwCF may have unexpected off-target effects, enhancing *Mabsc whiB7* expression and promoting *Mabsc* persistent infection.

**Significance Statement:** Pulmonary infections caused by *Mycobacterium abscessus* (*Mabsc*) are increasing in prevalence, especially in people with cystic fibrosis (pwCF) and bronchiectasis, and are very challenging to treat. Risk factors that predispose to *Mabsc* pulmonary infections remain poorly defined. This study demonstrates that tobramycin, an antibiotic commonly used to treat chronic *Pseudomonas aeruginosa* infections in pwCF, promotes *Mabsc* persistence by inducing *Mabsc* expression of the transcription factor *whiB7*. WhiB7 enhances *Mabsc* resistance to oxidative stress and survival in host macrophages. Transcriptome analysis reveals that tobramycin activates a large WhiB7 dependent regulon linked to stress tolerance and immune evasion.

These findings highlight a clinically relevant off-target effect of tobramycin that may contribute to *Mabsc* persistence in polymicrobial infections.

## Introduction

Airway infection with the non-tuberculous mycobacteria (NTM) species *Mycobacterium abscessus* complex (*Mabsc*) poses a major challenge for people with chronic airway disease, including bronchiectasis, and people with bronchiectasis caused by the genetic disease cystic fibrosis (CF) are at highest risk(1). *Mabsc* pulmonary disease (PD) is difficult to treat, and associated with poor clinical outcomes(1, 2). Cystic fibrosis transmembrane conductance regulator (CFTR) modulator therapies have greatly improved health in people with CF (pwCF), but do not appear to reverse bronchiectasis(3). As bronchiectasis is an independent risk factor for development of *Mabsc* lung disease(4), *Mabsc* infections are likely to remain a threat to pwCF. It thus remains imperative to determine what factors promote *Mabsc* persistence in order to identify those at risk and inform development of more effective and targeted therapies to eradicate *Mabsc* infections.

Prior studies have sought to identify risk factors for *Mabsc*-PD; however, beyond bronchiectasis being a prominent risk factor, studies have failed to detect commonalities among study cohorts(5–8). For example, while some find that chronic *P. aeruginosa* airway colonization is associated with *Mabsc* lung infection, others do not(7–10). Ultimately, susceptibility is likely multi-factorial, with different combinations of risk factors affecting different individuals.

As pwCF are both at highest risk for *Mabsc*-PD and a highly studied population(11), we looked to the CF literature for clues regarding potentially modifiable risk factors for *Mabsc* infection. We considered whether previously identified associations between CF airway co-infections and *Mabsc* lung infection could result from off-target effects of commonly used antibiotics. For instance, *Pseudomonas aeruginosa* infection is common in pwCF(11), and those with chronic *P. aeruginosa* infections are usually subjected to many courses of antibiotics including the aminoglycoside tobramycin(12, 13). Thus, if present, *Mabsc* is also exposed to these therapeutics. However, because *Mabsc* is inherently resistant to killing by tobramycin, it is generally assumed that administration of tobramycin affects neither viability nor fitness of *Mabsc*(14).

Recent studies have determined that sublethal doses of several ribosome targeting antibiotics used to treat *Mabsc* infections (the aminoglycoside amikacin; tetracyclines; and macrolide antibiotics, including azithromycin and clarithromycin) promote *Mabsc* antibiotic resistance by transcriptionally activating *whiB7* which encodes the transcription factor WhiB7(15– 20). WhiB7 is conserved in actinomycetes including *Mabsc, Mycobacterium smegmatis, Mycobacterium tuberculosis*, and *Streptomyces* spp., among others. In addition to playing a role in antibiotic resistance, across species WhiB7 functions as a global regulator, inducing genes involved in cell division and resistance to host stresses such as amino acid starvation, oxidative stress, and phagocytosis(20–22). Although studies have evaluated the crucial role of WhiB7 in resistance to antibiotics used to treat *Mabsc* infection, there is a notable deficiency in research exploring off-target effects of antibiotics aimed at other CF pathogens, such as tobramycin. Because of the frequency of co-infections of these two species (9), off-target effects of anti-pseudomonad treatment are a potential component of the complex dynamic between *Mabsc* and *P. aeruginosa* co-infections. Our study aimed to evaluate how tobramycin contributes to the risk of acquiring chronic *Mabsc* infections. Herein we demonstrate that administration of tobramycin to treat airway co-infections such as *P. aeruginosa* may inadvertently promote *Mabsc* persistence by inducing *whiB7* expression.

## Results

### Tobramycin induces *whiB7* gene expression in *Mabsc*

Clinically relevant concentrations of antibiotics frequently used in pwCF to kill bacteria other than *Mabsc* were evaluated for their ability to induce *whiB7* expression in *Mabsc*(12, 23, 24). Clinically relevant antibiotic concentrations were defined in two ways. First, we followed the Clinical Laboratory Standard Institute (CLSI) antibiotic breakpoints for *P. aeruginosa*(25, 26). Second, the peak concentration of antibiotic in the blood (C_max_) at clinically administered doses was considered (Table S1)(27–33). Based on these values the killing capacities of amikacin, tobramycin, ciprofloxacin, levofloxacin, and azithromycin were measured against our *P. aeruginosa* reference strain (Table S2). To ensure that antibiotics used in our studies would not kill *Mabsc*, minimum inhibitory concentration (MIC) assays against *Mabsc* reference strains ATCC 19977 and 390S(34) were performed (Figure 1a).

**Figure 1.**
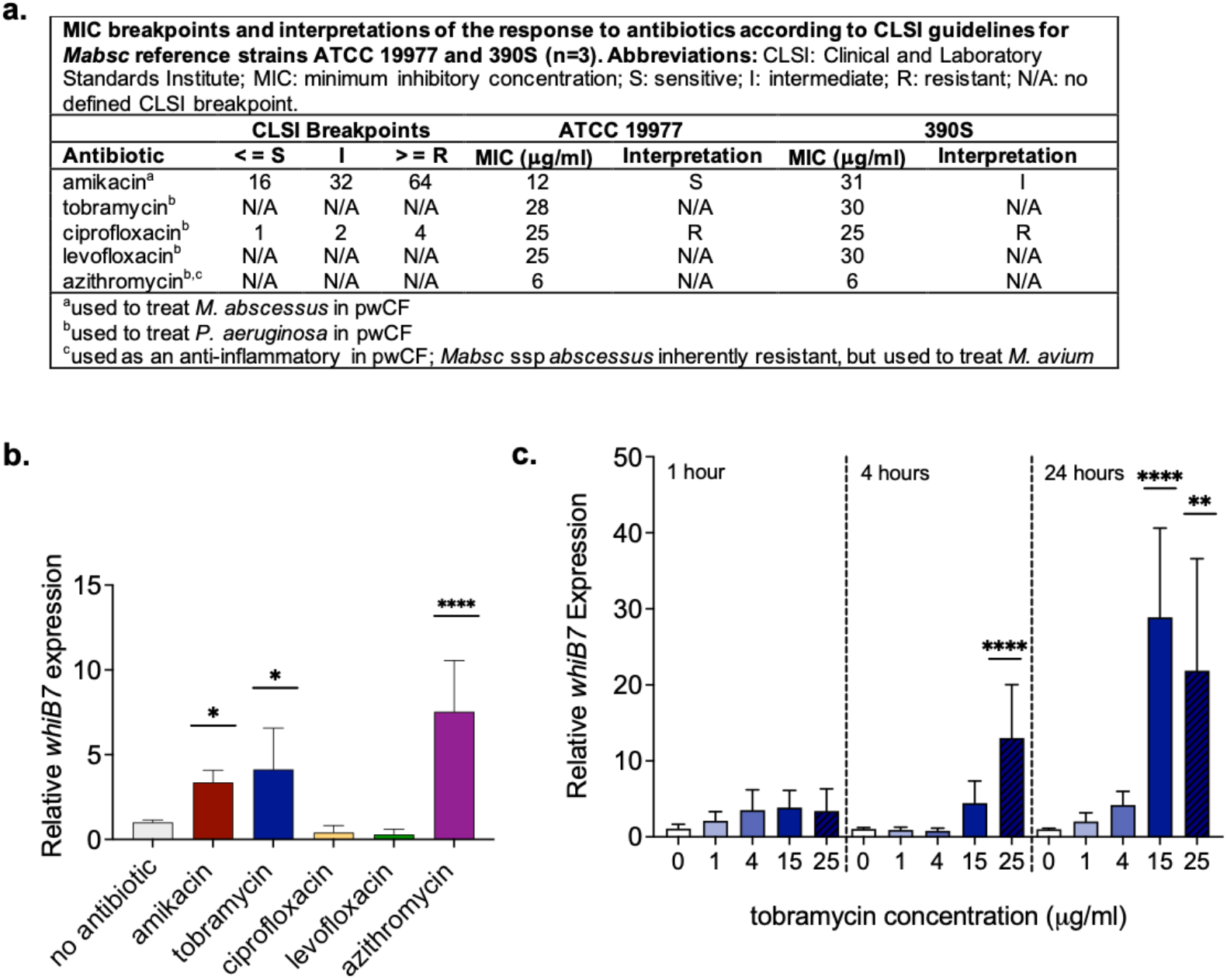
*Mabsc whiB7* expression is induced by tobramycin. **a)** Table depicting amikacin, tobramycin, ciprofloxacin, levofloxacin and azithromycin MIC breakpoints as defined by CLSI guidelines and as determined by experiment for *Mabsc* ATCC 19977 and 390S WT strains. **(b, c)** *Mabsc* ATCC 19977 *whiB7* transcripts were measured by qRT-PCR after *Mabsc* treatment with **b)** 1 μg/ml amikacin (positive control), 4 μg/ml tobramycin, 4 μg/ml ciprofloxacin, 4 μg/ml levofloxacin, or 5 μg/ml azithromycin for 24 hours or after **c)** treatment with 1, 4, 15, or 25 μg/ml tobramycin compared to no treatment at 1, 4, and 24 hours post tobramycin exposure. Statistical analyses for all experiments were performed by one-way ANOVA with Dunnett’s multiple comparisons post-hoc test relative to untreated WT at the same time point; *p <0.05, **p0.002, ****p < 0.0001, comparing each condition to no treatment at the same time point. Each bar represents the average of at least 3 experiments.

To test whether selected antibiotics induce *whiB7* in *Mabsc*, cultures of the *Mabsc* reference strain ATCC 19977 were grown in media containing tobramycin (15 µg/ml) or in sublethal doses of the control antibiotics amikacin (1 µg/ml) or azithromycin (4 µg/ml), ciprofloxacin (4 µg/ml) or levofloxacin (4 µg/ml) for 24 hours. Consistent with our hypothesis, tobramycin significantly induced *Mabsc whiB7* expression as compared to *Mabsc* cultures not exposed to antibiotics (Figure 1b, c). Induction of *whib7* expression was both dose- and time-dependent (Figure 1c). In contrast, fluoroquinolone antibiotics ciprofloxacin and levofloxacin, which are commonly used to treat *P. aeruginosa* infections in pwCF, did not induce *whiB7* gene expression at any tested concentration over 24 hours (Figure 1b, S1-S3). Induction of *whiB7* expression also occurred in the clinical isolate *Mabsc* 390S following exposure to amikacin and tobramycin (Figure S1). Since *Mabsc whiB7* expression plateaued at tobramycin doses greater than 15 µg/ml, and this concentration does not inhibit *Mabsc* growth, 15 μg/ml tobramycin was used to induce WhiB7 in subsequent experiments.

### Tobramycin enhances *Mabsc* resistance to reactive oxygen species and survival in macrophages

We next investigated how tobramycin induction of *Mabsc whiB7* expression could alter host-pathogen interactions. RNAseq studies have shown that the *Mabsc* WhiB7 regulon includes genes predicted to be important for survival in oxidative stress environments(20–22), such as the highly inflamed CF airway(35). We hypothesized that induction of *whiB7* expression by tobramycin would promote *Mabsc* resistance to reactive oxygen species (ROS). We investigated *Mabsc*’s resistance to oxidative stress by determining the hydrogen peroxide (H_2_O_2_) minimum inhibitory concentration (MIC) of *Mabsc* cultures. Pre-treatment of *Mabsc* with amikacin or tobramycin resulted in higher H_2_O_2_ MICs compared to untreated *Mabsc*, suggesting that both antibiotics may promote resistance to ROS in vivo (Figure 2a).

**Figure 2.**
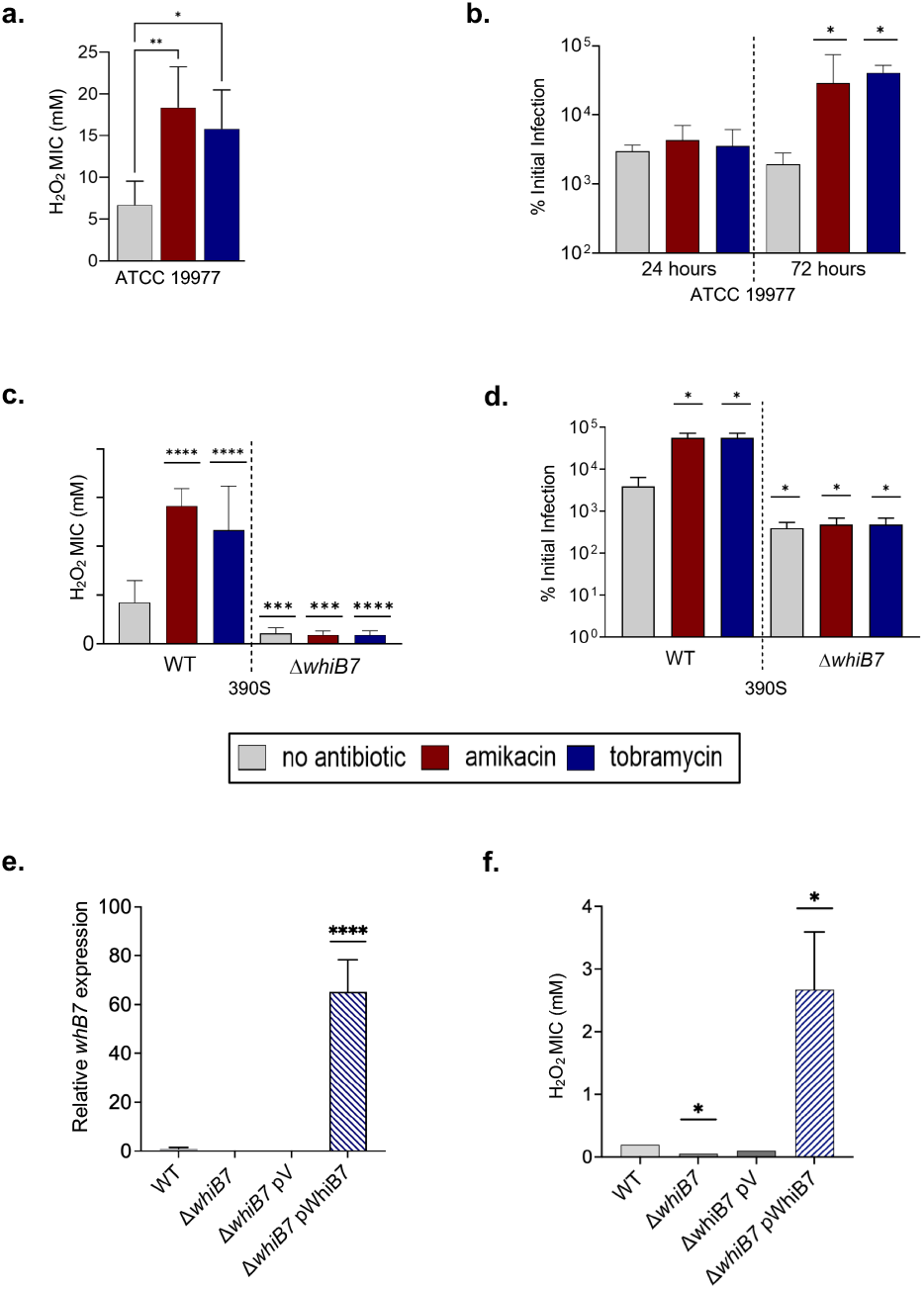
Tobramycin promotes *Mabsc* resistance to ROS and growth in human primary macrophages in a *whiB7*-dependent manner. **a)** *Mabsc* ATCC 19977 treated with 15 µg/mL tobramycin or 1 µg/mL amikacin exhibited increased resistance to H_2_O_2_, as determined by minimum inhibitory concentration (MIC) assays. **b)** Tobramycin and amikacin treated *Mabsc* showed enhanced intracellular survival in human monocyte-derived macrophages (MDMs) at 72 hours post infection compared to untreated *Mabsc*. **(c, d)** *Mabsc* 390S WT and Δ*whiB7* strains were treated with tobramycin or amikacin followed by assessment of **c)** H_2_O_2_ MICs or d) survival in MDMs. **(e, f)** Δ*whiB7* strains containing a plasmid constitutively expressing *whiB7* (pWhiB7) or empty vector (pV) were assessed for **e)** *whiB7* expression by qRT-PCR and **f)** H_2_O_2_ MICs. Statistical analyses for all experiments were performed by one-way ANOVA with Dunnett’s multiple comparisons post-hoc test relative to untreated WT; *p < 0.05, ***p<0.005, ****p<0.0001. MIC values are known between experiments. Each bar represents the average of at least 3 experiments.

RNAseq studies have also shown that the *Mabsc* WhiB7 regulon includes genes potentially important for invasion of macrophages, and *whiB7* is highly induced in *M. tuberculosis* after ingestion by macrophages(36, 37). Macrophages in the CF airways provide host defense against NTM species(38–40). As most macrophages found in the CF airway are monocyte derived macrophages (MDMs) recruited from the blood(38), evaluated the ability of *Mabsc* pretreated with aminoglycosides for ability to survive in MDMs. *Mabsc* exposed for 24 hours to either sublethal doses of amikacin or clinically relevant doses of tobramycin exhibited enhanced growth in human primary MDMs at 72 hours compared to untreated *Mabsc* (Figure 2b).

### The *Mabsc ΔwhiB7* mutant demonstrates impaired ROS resistance and growth in macrophages

To determine if enhanced resistance to host anti-microbial mechanisms following pre-treatment of *Mabsc* with tobramycin is dependent on induction of *whiB7*, we performed H_2_O_2_ MIC assays and macrophage infection assays using an *Mabsc whiB7* deletion mutant (*ΔwhiB7*) made in the background of the *Mabsc* clinical isolate 390S(41). As seen with *Mabsc* 19977, wild type (WT) 390S had a higher H_2_O_2_ MIC when exposed to sublethal concentrations of amikacin or anti-pseudomonal concentrations of tobramycin (Figure 2c). *Mabsc* Δ*whiB7* was more sensitive to killing by H_2_O_2_, with a significantly lower MIC compared to WT (Figure 2c). Furthermore, exposure of *Mabsc* Δ*whiB7* to either amikacin or tobramycin for 24 hours prior to exposure to H_2_O_2_ did not enhance resistance to H_2_O_2_ (Figure 2c).

*Mabsc* Δ*whiB7* also showed decreased growth in human primary MDMs compared to WT after 72 hours of infection (Figure 2d). Additionally, growth of *Mabsc* Δ*whiB7* was not restored to WT levels in human primary MDMs after *Mabsc* Δ*whiB7* was primed for 24 hours with tobramycin or amikacin (Figure 2d). Collectively, these data demonstrate the importance of *whiB7* expression for *Mabsc* survival in host cells, and support that the mechanism by which aminoglycosides enhance *Mabsc* ROS tolerance and enhanced growth in macrophages occurs via a WhiB7-mediated pathway(s).

### Overexpression of *whiB7* leads to increased ROS resistance

To confirm that enhanced ROS sensitivity of *Mabsc* Δ*whiB7* was due to loss of *whiB7*, not unmarked mutations, we complemented *Mabsc* Δ*whiB7* by introducing a constitutively expressed *whiB7* gene under control of the Hsp60 promoter on a multicopy, replicating plasmid (pWhiB7). Using qRT-PCR we found that the empty vector did not affect *whiB7* expression (comparison of *Mabsc* WT and pV *whiB7)*, and the complemented mutant over-expressed *whiB7* (Figure 2e). The *Mabsc* Δ*whiB7* isolate containing pWhiB7 was not only restored for resistance to H_2_O_2_, but also exhibited increased H_2_O_2_ MIC, demonstrating that increases in *whiB7* expression correlate with increased ROS resistance. The vector alone (pV) did not affect H_2_O_2_ MIC in *Mabsc* Δ*whiB7*, with an MIC similar to vectorless Δ*whiB7* (Figure 2f).

### Parenteral tobramycin improves survival of *Mabsc* in murine lungs during acute infection

To determine if tobramycin enhances *Mabsc* survival in the lung, we employed an acute model of murine *Mabsc* pneumonia (Figure 3). C57BL/6J mice were injected intraperitoneally (IP) with tobramycin (150 mg/kg)(42) 1 day prior to oropharyngeal (OP) inoculation with 5×10^6^ CFU *Mabsc* 19977, and then every day during a 7-day infection. Mice administered tobramycin demonstrated increased *Mabsc* colony forming units (CFUs) in the lungs at 7 days post infection (dpi) as compared to vehicle-treated mice (Figure 3a, b). To confirm that IP tobramycin levels reached clinically relevant concentrations in murine lungs, a second group of mice was dosed with IP tobramycin or vehicle via the same protocol, but were infected OP with 5×10^6^ CFU *P. aeruginosa*. Lungs from *P. aeruginosa* infected mice were harvested 1 dpi for CFU enumeration (Figure 3a, c). Vehicle treated mice demonstrated a 2.4 log decrease in CFUs in the lungs with > 2 × 10^4^ CFUs remaining. However, no *P. aeruginosa* CFUs were detected in the lungs of tobramycin treated mice, confirming that a therapeutic dose of tobramycin had been achieved (Figure 3c). Thus, administration of tobramycin at doses used to treat *Pseudomonas* infections can enhance survival of *Mabsc* in the lungs during acute infection.

**Figure 3.**
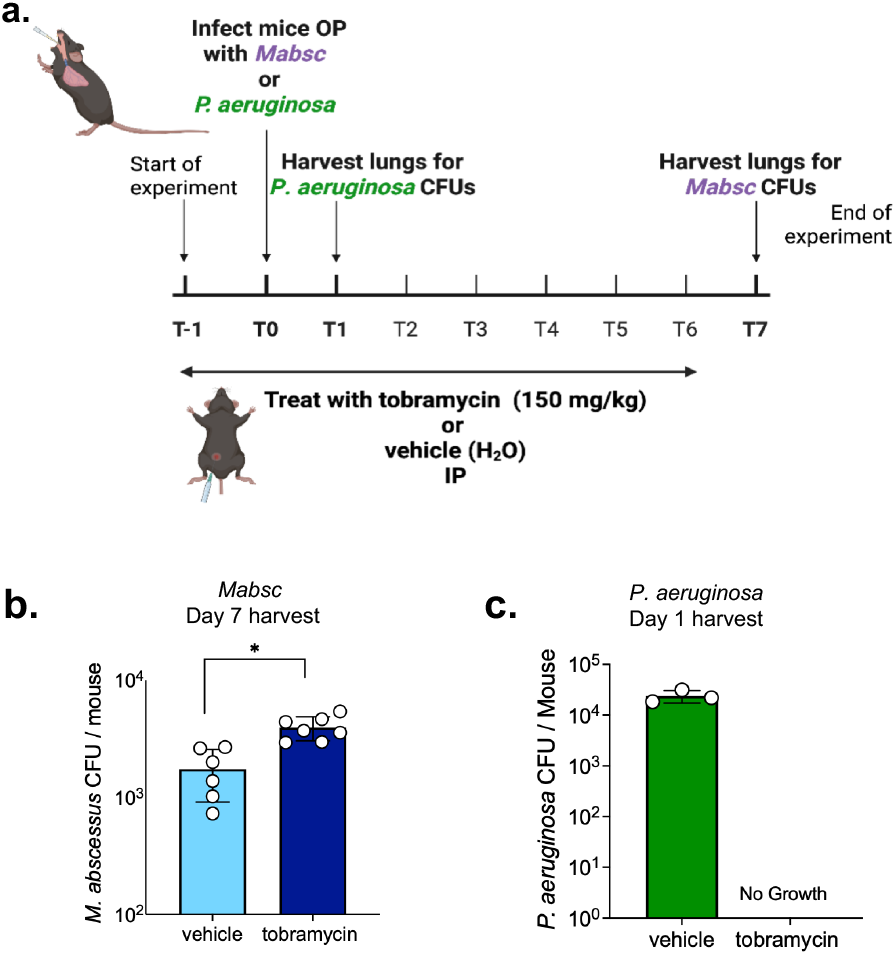
Administration of systemic tobramycin that achieves concentrations in murine lung sufficient to kill *P. aeruginosa* is associated with increased pulmonary *Mabsc* burden. **a)** Diagram of acute mouse model of *Mabsc* or *P. aeruginosa* infection for CFU quantification. Created with BioRender (https://www.biorender.com/). Mice were administered tobramycin (150 mg/kg, I.p.) or vehicle control 1 day prior to and on each day of pulmonary infection with *Mabsc* ATCC 19977 or *P. aeruginosa* via oropharyngeal aspiration. Lungs were harvested for CFU enumeration at 7 dpi *(Mabsc)* **(b)** or 1 dpi (*P. aeruginosa*) **(c)**. Statistical significance was determined by Student’s *t* test (* p < 0.05).

### RNA sequencing reveals high overlap between the tobramycin and WhiB7 dependent regulons

To characterize which components of the WhiB7 regulon were modulated by tobramycin, we compared gene expression profiles from *Mabsc* WT and Δ*whiB7* strains treated with tobramycin to untreated WT (Figure 4, Tables S3, S4,). Strikingly, we determined that of the approximated 5,000 annotated genes in the *Mabsc* genome, over half of the transcripts were differentially expressed (log_2_ fold change >1) when comparing tobramycin-treated to untreated (2742 genes), or when comparing the *Mabsc* Δ*whiB7* strain compared to *Mabsc* WT (2671 genes; Figure 4). In support of our hypothesis that tobramycin promotes expression of WhiB7, there were 2384 (80.68%) overlapping genes in the tobramycin-induced and WhiB7-dependent regulons, and expression of over 80% of overlapping genes were inversely regulated in tobramycin exposed *Mabsc* WT as compared to *Mabsc* Δ*whiB7* (Figure 4c, d). Additionally, only 149 genes were significantly changed in *Mabsc* Δ*whiB7* treated with tobramycin compared to *Mabsc* Δ*whiB7* not exposed to tobramycin (Figure 4c, d). All of the top 25 genes up- or down-regulated by tobramycin in *Mabsc* WT were inversely regulated in *Mabsc* Δ*whiB7* compared to WT, with only 7 genes also differentially expressed in *Mabsc* Δ*whiB7* treated with tobramycin compared to untreated *Mabsc* Δ*whiB7* (Tables S3, S4).

**Figure 4.**
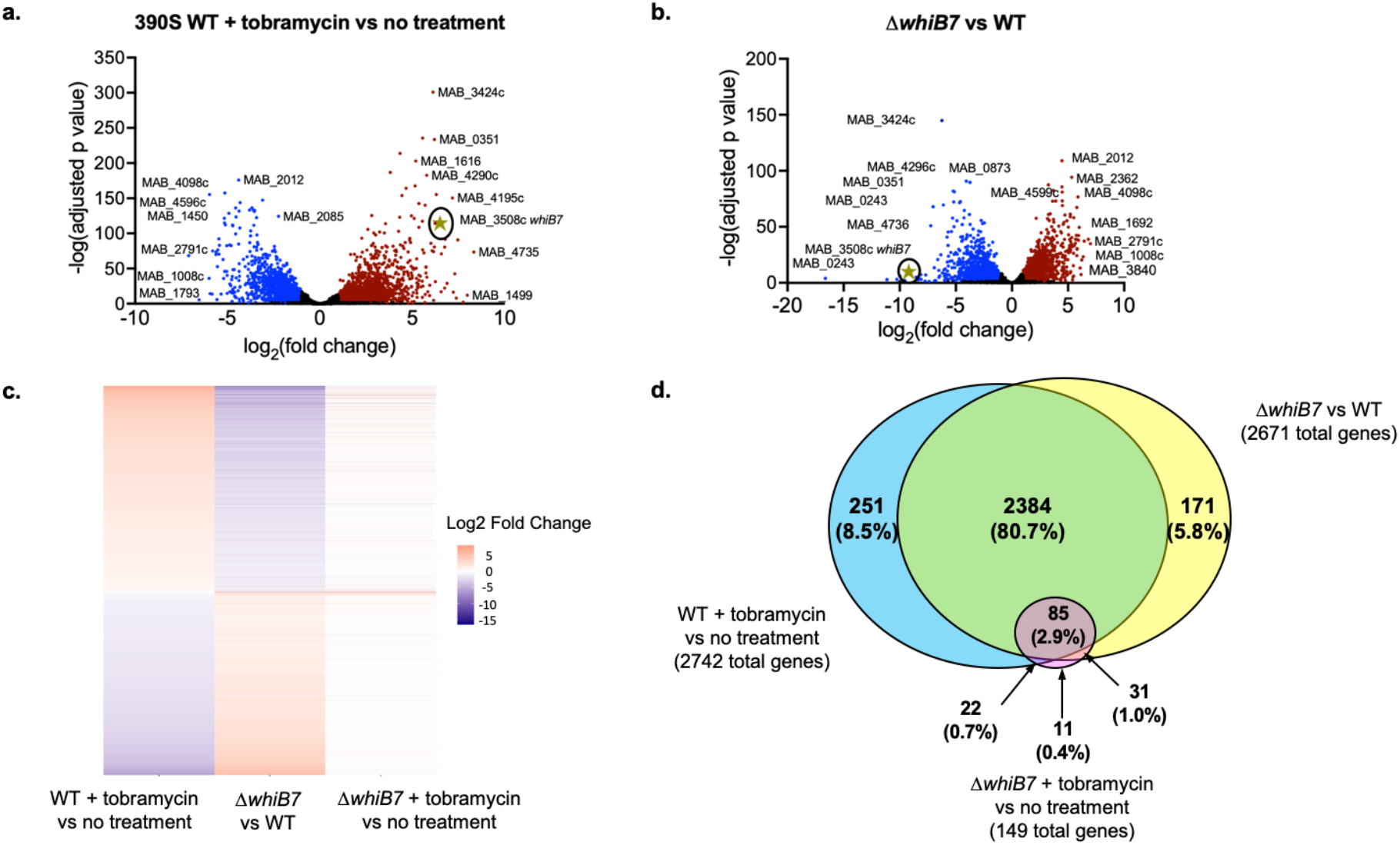
RNA-seq analysis of differential gene expression defining the tobramycin associated WhiB7 regulon. Volcano plots display differential expression profiles for **a)** *Mabsc* 390S WT treated with tobramycin (15 μg/ml) compared to untreated WT and **b)** 390S Δ*whiB7* compared to WT. Each gene is represented by a point; significantly up-regulated and down-regulated genes are shown in red and blue respectively and *whiB7* is represented by a circled gold star. **c)** Heat map of differentially expressed genes comparing the three conditions: WT treated with tobramycin compared to untreated WT, Δ*whiB7* compared to WT, and Δ*whiB7* treated with tobramycin compared to untreated Δ*whiB7*. **d)** Venn diagram summarizing the overlap of significantly differentially expressed genes among the three groups with number of genes and % of significant genes listed.

Consistent with the previously published WhiB7 RNAseq study(19), we identified several WhiB7 regulated genes responsive to tobramycin treatment that encode factors predicted to be important for macrophage survival and antibiotic resistance factors like *eis2* (MAB_4532c; S3, S4)(19, 43). The *Mabsc* WhiB7 and tobramycin regulons also included genes encoding factors that aid bacterial survival during oxidative stress conditions and phagocytosis, *katA* (MAB_0351)(37, 44) and *pknE* (MAB_4736)(45) as well as those that modulate components the *Mabsc* cell envelope, *fadE5* (MAB_4437)(46, 47) and *mps2* (MAB_4098c) (48, 49). Other WhiB7 and tobramycin regulated genes include those that elicit inflammatory toll like receptor (TLR)-2 responses in the host, such as *lpqH* (MAB_0885c; Tables S3, S4)(50). The large overlap in the *Mabsc* WhiB7 and tobramycin-responsive regulons further suggest that the mechanism by which tobramycin regulates the majority of its regulon occurs via modulating *whiB7* expression.

## Discussion

Although pwCF and bronchiectasis are at high risk of contracting *Mabsc*-PD, an unresolved question is why some pwCF develop persistent *Mabsc* infection, while others do not. Here we show that tobramycin, a standard of care aminoglycoside used to treat *P. aeruginosa* infections in pwCF(12, 13), upregulates *whiB7* and its downstream regulon(21), enhancing *Mabsc* resistance to oxidative stress and promoting survival in macrophages. These phenotypes were absent in a *whiB7* deletion mutant, emphasizing the importance of the WhiB7 regulon in shaping *Mabsc* physiology in ways that favor establishment of chronic infection.

WhiB7 is a conserved actinomycete transcription factor that connects antibiotic induced ribosomal stress to mycobacterial defense(18, 19, 51). Prior work has demonstrated that ribosome targeting antibiotics including aminoglycosides, macrolides, tetracyclines, and lincosamides induce *whiB7* expression(15–21), leading to upregulation of antibiotic resistance factors such as *eis2* and *erm(41)*(19). Extensive studies involving whiB7-mediated antibiotic resistance establish *whiB7* as a central hub of intrinsic drug resistance. Stimuli that induce *whiB7*, however, are not limited to antibiotics. Stressors that may be encountered in vivo can also induce *whiB7* (21, 52, 53). For example, *whiB7* is induced during *M. tuberculosis* infection in mouse lungs(53). We found that *Mabsc* exposure to doses of tobramycin that induce *whiB7* enhanced *Mabsc* survival in macrophages and in mice, indicating that aminoglycosides and other antibiotics may be more potent *whiB7* inducing stimuli than host factors.

Consistent with other reports, we found that ribosome targeting antibiotics (16, 21) induced *whiB7* expression; we add to these studies by demonstrating that fluoroquinolones ciprofloxacin and levofloxacin, which target bacterial DNA gyrase, did not induce *whiB7*. Previous RNAseq studies in *Mabsc* defined antibiotic associated WhiB7 regulons, including clindamycin (a lincosamide antibiotic that targets the ribosome)(54), and sub-inhibitory levels of amikacin(16), further demonstrating the importance of ribosome targeting antibiotics for induction of *whiB7*. Recent studies have proposed mechanisms by which ribosome stalling during translation, as would occur with ribosome targeting antibiotics, results in enhanced WhiB7 expression(55, 56). Thus, ribosomal stress activation by a variety of ribosome targeting antibiotics is a key mechanism for activation of *whiB7* expression and the subsequent phenotypes associated with induction of the WhiB7 regulon. However, to the best of our knowledge, this study is the first to specifically demonstrate that tobramycin, widely used to treat *P. aeruginosa* but not considered effective against *Mabsc*, can not only induce *whiB7*, but also program *Mabsc* to survive in vitro and in vivo stressors.

Apart from antibiotic resistance, WhiB7 mediates a broader range of adaptive stress responses that may promote immune evasion and host persistence. For example, *M. tuberculosis eis* (homolog of *Mabsc eis2*) suppresses macrophage activation and promotes intracellular bacterial survival by modulating host signaling pathways, including JNK pathway(57). Further, previous work demonstrated that amikacin treatment and overexpression of *whiB7* in *Mabsc* promoted survival in the macrophage-like THP-1 cell line(16). We have extended these data by showing that *whiB7* activation by tobramycin promotes *Mabsc* survival in primary human macrophages, and increasing levels of WhiB7 are associated with increasing resistance to reactive oxygen species.

A handful of studies have characterized the WhiB7 regulon across actinomycetes, revealing both conserved and species-specific genes(19–22, 51, 57, 58). RNAseq comparisons of the WhiB7 regulon in *Mabsc* and *M. smegmatis* revealed very limited overlap(19). Our transcriptomic profiling revealed that around half of the *Mabsc* genome was differentially expressed in response to either WhiB7 or tobramycin, with about 80% overlap between the two transcriptional responses. Shared WhiB7 and tobramycin induced genes include stress response genes, cell envelope regulators, and virulence-associated factors suggesting a convergent mechanism through which antibiotic exposure primes *Mabsc* for host adaptation. This is likely an evolutionary strategy for *Mabsc* to survive environmental stressors such as secreted antibiotics by soil microbes and predation by amoeba (59). Our gene expression threshold (log_2_ fold change >1) was more permissive than the prior Hurst-Hess et al RNAseq study defining the *Mabsc* WhiB7 regulon as 128 genes (log_2_ fold change >4)(19). We chose this more permissive threshold in order to capture physiologically relevant transcriptional changes. Most previously identified WhiB7 regulon genes were also detected in our studies (83 out of 128 genes). However, only 4 of the top 25 most regulated genes in the WhiB7 regulon identified in our studies and denoted in Table S3 were previously identified by Hurst-Hess. Furthermore, several key virulence determinants were identified in our study with log_2_ fold changes of just below the >4 cutoff. For example, *irtA* and *mmpS5*, genes important for encoding factors that regulate iron transport(60, 61), were found to be downregulated in our study only. The previous studies used the reference strain ATCC 19977 while ours used *Mabsc* 390S, suggesting that, besides different stringencies between our studies, strain to strain variability may account for differences observed in the WhiB7 regulon.

Several limitations of our study warrant discussion. While we demonstrated tobramycin-induced *whiB7* activity in two *Mabsc* strains, inter-strain variability in clinical isolates may influence the generalizability of *whiB7-*mediated responses. Future studies will assess *whiB7* induction and regulon activity in clinical *Mabsc* isolates from pwCF. We predict that differential tobramycin-induced expression of *whiB7* in clinical isolates may help explain the heterogeneity of susceptibility to *Mabsc-*PD in pwCF and *P. aeruginosa* infections. Additionally, our mouse infection model evaluates how tobramycin influences acute infection in healthy WT mice, a model that does not recapitulate the complex lung environment during chronic CF infection. Our ongoing work will explore how tobramycin and WhiB7 influence establishment of infection in murine models of chronic airway disease and during chronic *Mabsc* infection.

Even with these limitations, these findings have important clinical implications. First, our data highlight potential unintended consequences of standard antibiotic therapy in those with polymicrobial infections. Our observations support the concept that therapeutic strategies in pwCF and airways diseases should consider not only the direct antimicrobial effects of treatment, but also their potential off-target impacts on co-infecting bacteria. Prior studies in pwCF have noted unintended effects of antibiotics to enhance survival of non-targeted microbes. Azithromycin, which does not kill *Pseudomonas*, was shown to reduce the efficacy of inhaled tobramycin to kill *P. aeruginosa* in pwCF by inducing *P. aeruginosa* antibiotic resistance factors including MexXY (62). Off-target effects of antibiotics that enhance survival of bystander organisms likely occurs in other instances of polymicrobial infections beyond CF airways disease, such as other chronic lung diseases, chronic wounds, otitis media infections, and urinary tract infections(63, 64). Our results highlight that antibiotics can act as environmental cues that unexpectedly alter bacterial physiology and may enhance persistence of off-target organisms. Finally, our findings suggest that WhiB7’s dual role in antibiotic resistance and immune evasion may contribute to the challenge of eradicating *Mabsc* infections. Targeting global regulators like WhiB7 may offer new therapeutic opportunities. Understanding how antibiotic exposure shapes pathogen behavior and persistence will be critical for designing novel therapies for chronic infections in airways diseases like CF.

## Materials and Methods

### Bacterial strain and plasmid construction

Strains, plasmids, primer sequences, and cloning details are provided in supplementary Tables S5, S6. Routine cloning was performed with *E. coli* DH5α cultured in Luria Bertani (LB; Difco) medium with kanamycin (Research Products International) (50 μg/ml) as required. The *whiB7* expression plasmid was generated using primer pairs listed in Table S6 and cloned by isothermal assembly into pSD5.Hsp60(65). *Mabsc* Δ*whiB7* was constructed and donated by Kyle Rohde, PhD (University of South Florida)(41). *For additional details, see SI Appendix, SI Materials and Methods*.

### Antibiotic MIC assays

For *P. aeruginosa* MIC assays, 18-hour cultures of *P. aeruginosa* PA14 were centrifuged at 13,000xg for 1 minute, washed, resuspended in PBS to an A_600_ of 0.08 to 0.1 (1.5 × 10^8^ CFU), and then 100 μl of bacteria were added to 100 μl of Mueller Hinton Broth (Oxoid; MHB) in 96-well plates. Plates incubated statically for 24 hours at 37°C with humidity (Table S2). For *Mabsc* MICs, 3 day cultures of *Mabsc* strains ATCC 19977 and 390S WT were washed, then diluted to 2 × 10^6^ CFU/ml in 7H9-ADC-glycerol. Subsequently 100 μl of bacteria were added to 100 μl of 7H9-ADC-glycerol containing serial dilutions of amikacin, tobramycin, ciprofloxacin, levofloxacin, and azithromycin (concentrations of 0-100 μg/ml) in 96 well plates, then grown in a humidified incubator at 37°C for 5 days (Figure 1a). The lowest concentration of antibiotic inhibiting bacterial growth was defined as the MIC.

### Quantitative Reverse Transcription PCR (qRT-PCR)

RNA was extracted using the Monarch Total RNA Miniprep Kit (New England Biolabs) per the manufacturer instructions. cDNA generation and amplification were performed with PrimeTime qPCR primer and probe sets (Integrated DNA Technologies, Coralville, IA) listed in Table S6 in a one-step reaction with the Luna Probe One-Step RT-qPCR 4X Mix with UDG (New England Biolabs) per manufacturer’s protocol. Relative abundance of transcripts was quantified by determining the difference of target transcript compared to a reference gene (*rpoB*) by calculating 2^-ΔΔCT^ values. *For additional details, see SI Appendix, SI Materials and Methods*.

### Measurement of hydrogen peroxide MICs

*Mabsc* cultures were diluted to a cell density of 1 × 10^6^ CFU in PBS, to which varying concentrations (0-150mM) of 30% H_2_O_2_ (Fisher; 9.8M) were added. After 24 hours, 100 μl was plated to 7H10-OADC-glycerol plates to assess for growth. The lowest concentration of H_2_O_2_ inhibiting bacterial growth, as determined by no growth on plates, was defined as the MIC. Experiments were performed at least 3 times.

### Generation of monocyte derived macrophages (MDMs)

Blood was obtained from healthy human donors with IRB approval. Peripheral blood mononuclear cells (PBMCs) were isolated using density gradient centrifugation with Ficoll-Plaque Plus (Cytiva), and monocytes were purified from PBMCs via negative selection (Miltenyi Monocyte Isolation Kit II). Monocytes were plated at a density of 2.5 × 10^5^ cells per well in 48 well plates, and differentiated to MDMs using recombinant murine M-CSF as described(66). Monocyte and MDM culture media did not contain antibiotics to prevent confounding influences on infection assays.

### Macrophage infection models

*Mabsc* were added to MDMs at a multiplicity of infection (MOI) 1:1. MDMs were centrifuged for 1 minute at 1000xg to enhance microbe and macrophage contact, then incubated at 37 °C in 5% CO_2_. After 60 minutes incubation, MDMs were washed 3X in HBSS to remove extracellular bacteria, then 1) lysed with 0.1% triton (Sigma-Aldrich) in 1X PBS and serially diluted in PBS with 0.1% triton before plating on either LB agar for ATCC 19977 isolates or 7H10-OADC-glycerol for 390S isolates to determine *Mabsc* uptake by quantification of intracellular, or 2) fresh media were added and cells were incubated at 37 °C in 5% CO_2_ for 24 or 72 additional hours then washed and lysed with 0.1% triton in PBS as previously described. Experiments were performed at least 3 times.

### Mouse infection model

Male and female C57BL/6J (WT) mice, 8-9 weeks old were inoculated with *Mabsc* 19977 (5×10^6^ CFUs per mouse) or *P. aeruginosa* PA14 (5 × 10^6^ CFU per mouse) via oropharyngeal (o.p.) aspiration. One day prior to infection, mice were treated via intraperitoneal (i.p.) injection with tobramycin (150 mg/kg, ~ 150 μl per mouse of 20mg/ml tobramycin in H_2_O). Vehicle treated mice received 150 μl of water. At noted time points, mice were euthanized and lungs were removed for homogenization and plating for CFU quantification as described(66). *For additional details, see SI Materials and Methods*.

### RNA sequencing analyses

Late log cultures (3 days) of *Mabsc* 390S WT and Δ*whiB7* were back-diluted into 7H9-ADC-glycerol media with or without tobramycin (15 μg/ml) for 24 hours (3 replicates per condition), then processed for RNA as previously described using the Monarch Total RNA Extraction kit per manufacturer’s protocol. Ribosomal RNA was depleted using the NEBNext rRNA Depletion Kit (Bacteria; New England Biolabs) per manufacturer’s specifications. Quality control of RNA was performed using the TapeStation (Agilent). RNA library preparation, amplification, and sequencing were performed by the National Jewish Health Genomics Core. Library size (base pair), concentration, and quality scores were calculated via TapeStation. Libraries were sequenced using Illumina NextSeq 2000 at a sequencing depth of 50 million paired end reads per sample. *For additional details, see SI Appendix, SI Materials and Methods*.

## Statistical analyses

*For statistical analyses used in this study, see SI Appendix, SI Materials and Methods*.

## Acknowledgments

This work was supported by funding from the CF Foundation awards CORLEY23F0, HISERT19R3, NICK20Y2-SVC, NICK20Y2 and NIH award R01HL167956. We thank Dr. Kyle Rohde for supplying the *Mabsc* 390S WT and Δ*whiB7* strains, Dr. William DePas providing the pSD5.Hsp60 empty vector, the National Jewish Health Genomics Core for RNA sequencing and KC Anderson for RNAseq analysis, Dr. Michael Strong for providing *M. abscessus* gene annotations, and Dr. Elaine Epperson for assistance with RNA extraction protocols.

