## Supporting Information for "Tobramycin enhances *Mycobacterium abscessus* fitness through *whiB7* induction"

### Supporting Information for
#### Tobramycin enhances *Mycobacterium abscessus* fitness through *whiB7* induction

Jodi M. Corley[a*], Jack H. Congel[a], Kelsey C. Haist[a], Alma E. Ochoa[a], Kenneth C. Malcolm[a], William J. Janssen[a], Jerry A. Nick[a], and Katherine B. Hisert[a*]

*Corresponding authors: Jodi M. Corley


**This PDF file includes:**

SI Materials and Methods
Figures S1 to S3
Tables S1 to S6
SI References

**Supporting Information Materials and Methods.**

**General bacterial growth conditions.** *Pseudomonas aeruginosa* reference strain PA14, maintained as a stock frozen at -80°C, was grown in LB broth shaking (200 rpm) for 18 hours at 37°C prior to experiments. *Mycobacterium abscessus* ssp. *abscessus* strains ATCC 19977 smooth morphotype and 390S (also smooth morphotype), maintained as -80°C frozen stocks, were grown in Middlebrook 7H9 broth (Sigma-Aldrich) base with 10% Middlebrook Albumin Dextrose Catalase (ADC; Sigma-Aldrich) and 0.5% glycerol (Fisher; 7H9-ADC-glycerol), or on Middlebrook 7H10 agar (Difco) with 10% Oleic Acid Albumin Dextrose Catalase (OADC; Remel) and 0.5% glycerol (7H10-OADC-glycerol) or Luria-Bertani (LB) agar (Difco). Prior to each experiment, *Mabsc* cultures grown for 3 days at 37°C with shaking incubation (200 rpm) were centrifuged at 13,000xg for 2 minutes, washed with Phosphate Buffered Saline pH 7.4 (PBS; Gibco), then diluted to an $A_{600}$ of 0.08 to 0.1 ($1.5 \times 10^8$ CFU) in 7H9-ADC-glycerol with or without antibiotic (antibiotic identity and dose listed in Table 1), then grown at 37°C for 24 hours unless otherwise specified. Maintenance of ATCC 19977 and 390S WT isolates each containing a plasmid were grown in 7H9-ADC-glycerol with 50 μg/ml kanamycin and the 390S *whiB7* deletion strain (Δ*whiB7*) with a plasmid was grown in 7H9-ADC-glycerol and 250 μg/ml kanamycin as required.

**Gene expression studies; Quantitative Reverse Transcription PCR (qRT-PCR).** *Mabsc* cultures were centrifuged at 1200Xg for 20 minutes, supernatants were removed and pellets were resuspended Tris-EDTA (TE; Sigma) buffer, then centrifuged at 15000xg for 10 minutes. Pellets were resuspended in TE containing 2 mg lysozyme (chicken egg white; Thermo Scientific) and 0.005 mg Proteinase K (Invitrogen) then incubated at room temperature (RT) for 30 minutes(1). RNA was extracted using the Monarch Total RNA Miniprep Kit (New England Biolabs) per the manufacturer instructions. cDNA generation

and amplification were performed with PrimeTime qPCR primer and probe sets (Integrated DNA Technologies, Coralville, IA). Briefly, reactions contained 100 ng of RNA, 1 µL oligo/probe mix (0.4 $\mu$M of specific primer; 0.2 $\mu$M probe), 2.5 µL Luna Probe One-Step RT-qPCR 4X mix with UDG (New England Biolabs), 1 U RNase inhibitor RNase Out (Invitrogen), and RNase free water to a total volume of 10 µL per reaction. Reverse transcription and qPCR were performed and transcripts measured using a Bio-Rad thermal cycler, with the following cycle conditions: carryover prevention at 25°C for 30 seconds; reverse transcription at 55°C for 10 minutes; initial denaturation at 95°C for 1 minute; 35 cycles of denaturation at 95°C for 10 seconds and extension at 60°C for 30 seconds. Relative abundance of transcripts was quantified by determining the difference of target transcript compared to a reference gene (*rpoB*) by
calculating $2^{-\Delta\Delta CT}$ values. Primers and Probes are listed in Table S3.

**Mouse Infection Model.** Male and female C57BL/6J (WT) mice, 8-9 weeks old, were purchased from Jackson Labs. To prepare infection inoculum: *Mabsc* ATCC 19977 or *P. aeruginosa* PA14 was centrifuged at 14,000xg for 1 minute, washed with PBS twice, resuspended in PBS, and diluted to 1 x $10^8$ CFUs in 50 µl of PBS. Mice were inoculated with 50 µl of *Mabsc* (5x$10^6$ CFUs per mouse) or 50 µl of *P. aeruginosa* (5 x $10^6$ CFU per mouse) via oropharyngeal (o.p) aspiration. The murine pulmonary infection model was performed two times, with 4 or 5 mice per group in each experiment. O.p. inoculation was performed as described(2). One day prior to infection, mice were treated via intraperitoneal (i.p.) injection with tobramycin (150 mg/kg, ~ 150 µl per mouse of 20mg/ml concentration tobramycin in $H_2O$). Vehicle treated mice received 150 µl of water. Mice were euthanized as specified time points via i.p. injection of phenobarbital solution (Fatal-Plus; Vortech Pharmaceuticals, LTD). Lungs were removed for homogenization and plating for CFU quantification as described(2). All experiments using animals were approved by the Institutional Animal Care and Use Committee (IACUC) (Protocol number:

AS2574-02-23) at National Jewish Health. Mice were anesthetized with isoflurane prior to o.p. instillations. Following treatment, mice were examined daily for signs of distress or imminent death.

**RNA sequencing analyses.** Quality control and processing of the reads to gene-level abundances was performed using the nf-core RNA-seq pipeline (version 3.9)(3) for the Nexflow workflow tool (version 22.04.0)(4). Illumina universal adapters were removed from the reads with TrimGalore (version 0.6.7)(5). The quality of the reads was assessed using FastQC (version 0.11.9)(6) [Andrews, 2010] before and after read trimming. Reads were mapped with the STAR aligner (version 2.7.10a)(5) to the *Mabsc* ATCC 19977 genome assembly using gene annotations provided from NCBI (https://www.ncbi.nlm.nih.gov/datasets/genome/GCF_000069185.1). Ribosomal sequences were filtered out using the "--remove_ribo_rna" pipeline flag. Transcripts were quantified with Salmon (version 1.5.2)(7), and gene-level abundances were provided with the tximeta package (version 1.8.0)(8). Statistical comparisons were performed with DESeq2 (version 1.44.0)(9) on the bias-corrected gene counts from Salmon using Wald tests for group comparisons.

Analyzed RNAseq data was represented using volcano plots generated on GraphPad Prism 10 (GraphPad Software, Boston, MA; Figure 4a,b) and plotted based on $\log_2$. Venn diagrams were adapted from diagrams generated using the Venny2.1 software(10) (Figure 4c.). Heatmaps were generated in RStudio(11) (v2025.05.1+513) using the ggplot2 package (v3.5.1)(12). Differentially expressed genes were selected based on an adjusted $\log_2$ fold change >1 and -log(adjusted p-value). Data were plotted into long format using the tidyr(13) and dplyr packages(14), and the heatmap was constructed using geom_title. geom_tile(). Rows (genes) and columns (*Mabsc* samples and conditions) were optionally ordered based on hierarchical clustering. Expression values were scaled by gene to highlight relative differences across samples. Color gradients were applied using scale_fill_gradient2(ow="navy", mid="white", high="red", midpoint=0) (Figure 4d)(12).

**Statistical analyses.** Data analyses comparing two groups of normal distribution were analyzed by two-tailed unpaired Student's *t-* tests, p-value < 0.05. Normality of data was assessed using the Shapiro-Wilk test, p-value < 0.05. One-way analysis of variance with Tukey's posttest were used to determine statistical significance of three or more groups, p-value < 0.05. Statistical analyses and graphs were created using Prism 10 (GraphPad Software, Boston, MA) or R studio.

a.

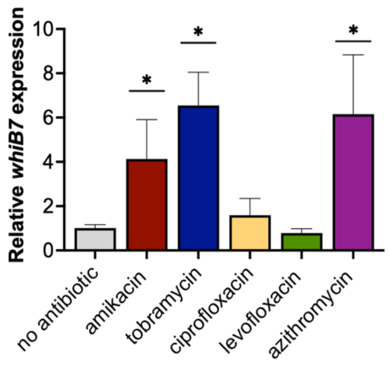

b.

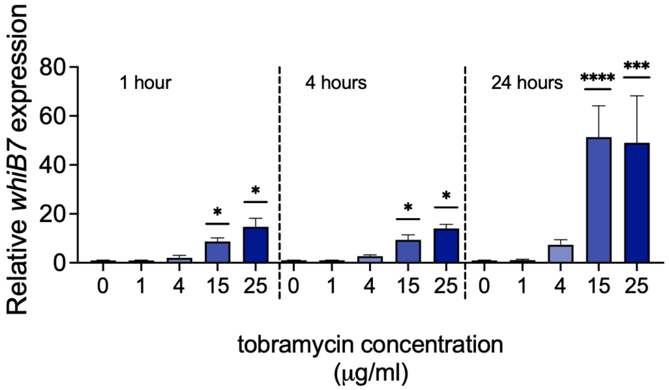

**Fig. S1. *Mabsc* 390S *whiB7* expression is induced by tobramycin.** *Mabsc* 390S *whiB7* transcripts were measured by qRT-PCR after *Mabsc* treatment with **a)** 1 µg/ml amikacin (positive control), 4 µg/ml tobramycin, 4 µg/ml ciprofloxacin, 4 µg/ml levofloxacin, or 5 µg/ml azithromycin for 24 hours or after **b)** treatment with 1, 4, 15, or 25 µg/ml tobramycin compared to no treatment at 1, 4, and 24 hours post tobramycin exposure. Statistical significance was determined by one-way ANOVA, *$p < 0.05$, ***$p < 0.0002$, ****$p < 0.0001$ as compared to no antibiotic treatment at the same time point.

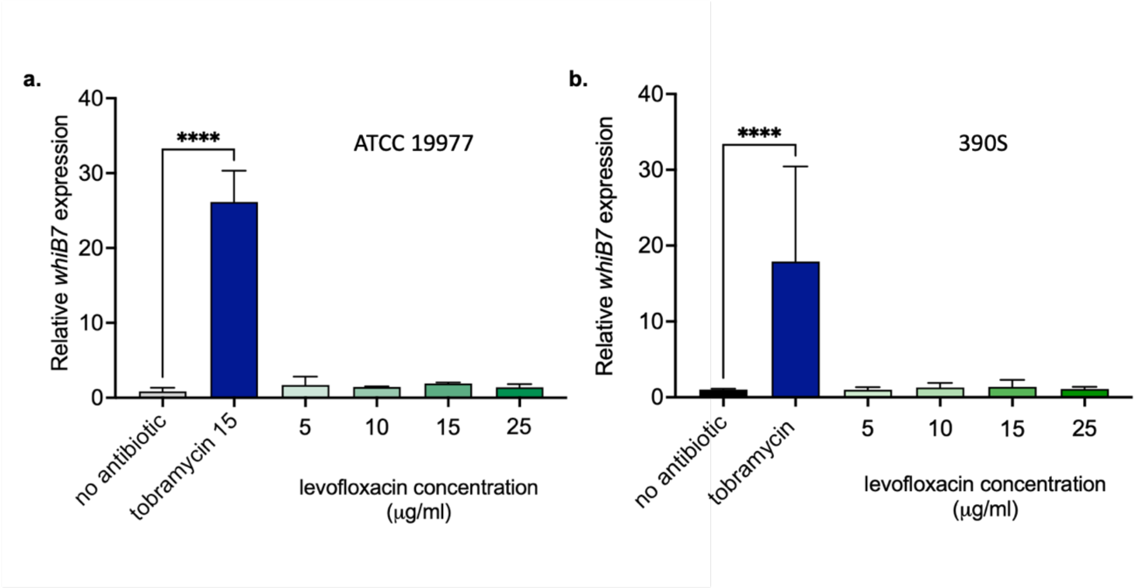

**Fig. S2. Levofloxacin does not induce *Mabsc whiB7* expression.** *Mabsc* **a)** ATCC 19977 and **b)** 390S *whiB7* transcripts were measured by qRT-PCR after *Mabsc* treatment with 5, 10, 15, or 25 μg/ml levofloxacin and compared to no treatment or the positive control tobramycin (15 μg/ml). Statistical significance was determined by comparing each condition to no antibiotic via one-way ANOVA with Dunnett's multiple comparisons post-hoc test; ****$p < 0.0001$.

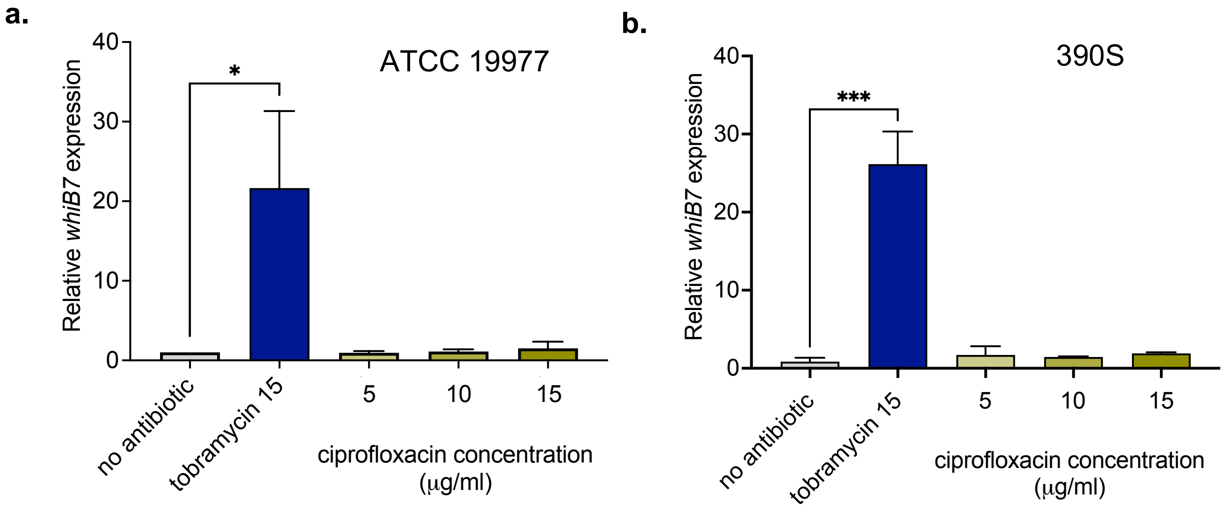

**Figure S3. Ciprofloxacin does not induce *Mabsc whiB7* expression.** *Mabsc* **a)** ATCC19977 *WT* and **b)** 390S WT*whiB7* transcripts were measured by qRT-PCR after *Mabsc* treatment with 5, 10, 15, or 25 µg/ml ciprofloxacin and compared to no treatment or the positive control tobramycin (15 µg/ml). Statistical significance was determined by comparing each condition to no antibiotic via one-way ANOVA with Dunnett's multiple comparisons post-hoc test; *p < 0.05, ***p < 0.0002.

**Table S1. Identity and characteristics of antibiotics used in this study.**

| Antibiotic | Microbial Target in CF | Class | Mechanism of action | Manufacturer | Reference |
|---|---|---|---|---|---|
| amikacin | *Mabsc* | aminoglycoside | Inhibits protein synthesis | Research Products International (RPI) | (15) |
| tobramycin | *P. aeruginosa* | aminoglycoside | Inhibits protein synthesis | In vitro: RPI<br>In vivo (pharmaceutical grade): Biosynth | (16) |
| ciprofloxacin | *P. aeruginosa* | Fluoroquinolone | Inhibits DNA synthesis | Thermo | (17) |
| levofloxacin | *P. aeruginosa* | Fluoroquinolone | Inhibits DNA synthesis | Thermo | (17) |
| azithromycin | *Mabsc* | Macrolide | Inhibits protein synthesis | Tocris Bioscience | (18) |

**Table S2. MIC breakpoints and interpretations of *P. aeruginosa* reference strain PA14 response to antibiotics according to CLSI guidelines. Abbreviations:** CLSI: Clinical and Laboratory Standards Institute; MIC: minimum inhibitory concentration; S: sensitive; I: intermediate; R: resistant.

| | **CLSI Breakpoints** | | | | **PA14** | |
|---|---|---|---|---|---|---|
| **Antibiotic** | **< = S** | **I** | **> = R** | $\mathbf{C_{max}}$ | **MIC (μg/ml)** | **Interpretation** |
| amikacin[a] | 16 | 32 | 64 | 64–80 μg/ml | 2 | S |
| tobramycin[b] | 1 | 2 | 4 | 20-30 μg/mL | 1.5 | I |
| ciprofloxacin[b] | 0.5 | 1 | 2 | 2.84 – 3.8 μg/ml | 0.19 | S |
| levofloxacin[b] | 1 | 2 | 4 | 2.8 - 5.2 μg/mL | 0.25 | S |
| azithromycin[c] | N/A | N/A | N/A | 0.6-2.1 μg/ml | >256 | N/A |

[a]used to treat *M. abscessus* in pwCF
[b]used to treat *P. aeruginosa* in pwCF
[c]used as an anti-inflammatory in pwCF; *Mabsc* ssp *abscessus* inherently resistant, but used to treat *M avium*

**Table S3.** Top 25 up regulated genes in *Mabsc* treated with tobramycin vs untreated with expression compared to: Δ*whiB7* vs WT and Δ*whiB7* + tobramcyin vs no treatment, as determined by RNAseq.

| | | | Log$_2$(fold Δ) | | |
|---|---|---|---|---|---|
| **Gene locus** | **Gene name** | **Product or function** | **WT + tobramycin vs untreated** | **Δ*whiB7* vs WT** | **ΔwhiB7 + tobramycin vs untreated** |
| MAB_4735[#] | *dps2* | Putative starvation-induced DNA protecting protein/Ferritin | 8.33238779 | -6.60748119 | 1.724907 |
| MAB_1499 | *dadA1* | Putative FAD dependent oxidoreductase | 7.976316527 | -8.122916225 | -0.1466* |
| MAB_0243 | | Hypothetical_protein | 7.749520457 | -16.63571698 | -8.8862 |
| MAB_0734 | | MspA_family_protein | 7.462791362 | -5.330935661 | 2.131856 |
| MAB_0046 | *PE5_1* | PE family immunomodulator PE5 | 7.378655904 | -5.646852576 | 1.731803** |
| MAB_2386 | | Conserved hypothetical protein | 7.298058142 | -4.661479811 | 2.636578 |
| MAB_2769 | *cyoC* | Cytochrome bo(3) ubiquinol oxidase subunit 3 | 7.220359538 | -6.024309116 | 1.19605** |
| MAB_4195c | *mdtD2* | Putative multidrug resistance protein MdtD | 7.170687707 | -7.020387355 | 0.1503* |
| MAB_0461 | | hypothetical_protein | 6.873952321 | -6.509083318 | 0.364869* |
| MAB_4736 | *pknE* | Serine/threonine-protein kinase PknE | 6.778673119 | -7.227098057 | -0.44842* |
| MAB_4268c | | Hypothetical protein | 6.753416014 | -5.596907655 | 1.156508** |
| MAB_1264 | | Hypothetical protein | 6.749056309 | -3.071748236 | 3.677308 |
| MAB_0735 | | Conserved hypothetical protein | 6.573446991 | -6.328980888 | 0.244466* |
| MAB_1321 | | MspA_family_protein | 6.559604052 | -5.296384274 | 1.26322** |
| MAB_3508c[#] | *whiB7* | Transcriptional regulator | 6.50107715 | -9.199352573 | -2.69828** |
| MAB_4295c | *udgA/rkpK* | UDP-glucose 6-dehydrogenase | 6.299155001 | -6.111842662 | 0.187312 |
| MAB_0472 | | Type II secretion system protein | 6.249963755 | -5.665201234 | 0.584763* |
| MAB_2740c | | Probable oxidoreductase | 6.229595673 | -5.680789475 | 0.548806* |
| MAB_0351 | *katA* | Catalase | 6.197640348 | -5.213821046 | 0.983819* |
| MAB_0540 | | Exopolyphosphatase | 6.196533014 | -5.003173774 | 1.193359 |
| MAB_0288 | *xlnD* | 3-hydroxybenzoate 6-hydroxylase | 6.154038231 | -9.968775916 | -3.81474** |
| MAB_3340[#] | *sasA2* | Probable sensor histidine kinase | 6.153377771 | -3.943097567 | 2.21028 |
| MAB_3424c[#] | | 2-isopropylmalate synthase | 6.124361129 | -6.236869704 | -0.11251* |
| MAB_2313 | | Conserved hypothetical protein | 6.068759169 | -5.697787306 | 0.370972* |
| MAB_1156c | *lysA1* | Probable diaminopimelate decarboxylase LysA | 6.011115058 | -5.007629306 | 1.003486** |

[#] Identified as part of the WhiB7 regulon previously by Hurst-Hess et al. 2017(19)
* Not statistically significant
**Not statistically significant based on p-adj value

**Table S4.** Top 25 down regulated genes in *Mabsc* treated with tobramycin vs untreated with expression compared to: Δ*whiB7* vs WT and Δ*whiB7* + tobramcyin vs no treatment, as determined by RNAseq.

| | | | Log$_2$(fold Δ) | | |
|---|---|---|---|---|---|
| **Gene locus** | **Gene name** | **Product or function** | **WT + tobramycin vs untreated** | **Δ*whiB7* vs WT** | **Δ*whiB7* + tobramycin vs untreated** |
| MAB_2791c | | Ethanolamine permease | -7.107304408 | 7.011941252 | -0.09536* |
| MAB_1793 | | Phage associated protein | -6.543335084 | 5.412140933 | -1.13119** |
| MAB_1008c | | MCE family protein Mce6C | -5.999303821 | 6.350522152 | 0.351218 |
| MAB_4098c | *mps2* | Peptide synthetase | -5.979564451 | 5.887673081 | -0.09189* |
| MAB_4354 | *rutE* | Putative malonic semialdehyde reductase | -5.943028289 | 5.51210953 | -0.43092* |
| MAB_4355 | *yisK_3* | Hypothetical fumarylacetoacetate (FAA) hydrolase family | -5.838424229 | 6.165251058 | 0.326827* |
| MAB_1513 | *acpS* | Holo-[acyl-carrier-protein] synthase | -5.806749289 | 5.944565951 | 0.137817* |
| MAB_1452 | *atpG* | ATP synthase gamma chain | -5.622828316 | 5.156214013 | -0.46661* |
| MAB_1451 | *atpA* | ATP synthase subunit alpha | -5.620709399 | 5.383549545 | -0.23716* |
| | *whiB_2* | | -5.570738402 | 5.444126794 | -0.12661* |
| MAB_2790c | *eutB_1* | Ethanolamine ammonia-lyase large subunit | -5.551729663 | 5.491182609 | -0.06055* |
| MAB_1453 | *atpD* | ATP synthase subunit beta | -5.527739825 | 5.140736341 | -0.387* |
| MAB_4105c | *mtfD* | Methyltransferase | -5.470555203 | 5.334356365 | -0.1362* |
| MAB_4106c | *atf1* | Acetyltransferase | -5.465820806 | 5.516115136 | 0.050294* |
| MAB_1454 | *atpC* | ATP synthase epsilon chain | -5.439045053 | 5.155883357 | -0.28316* |
| MAB_2609 | | Possible fatty-acid-CoA ligase FadD | -5.353142586 | 4.431395475 | -0.92175* |
| MAB_1884c | *aceE* | Pyruvate dehydrogenase E1 component | -5.249492169 | 4.96096427 | -0.28853* |
| MAB_1009c | | Mce family protein Mce5B | -5.215901154 | 5.79401661 | 0.578115* |
| MAB_1450 | *atpFH* | ATP synthase subunit delta | -5.181193465 | 4.909826828 | -0.27137* |
| MAB_1933c | *glnA_2* | Glutamine synthetase | -5.167456871 | 5.206373471 | 0.038917* |
| MAB_4815 | | Conserved hypothetical protein | -5.154123644 | 4.782096929 | -0.37203* |
| MAB_2362 | | Thiamin pyrophosphokinase | -5.149822157 | 5.32567965 | 0.175857* |
| MAB_4107c | *gtf1* | glycosyltransferase | -5.11828739 | 5.20748341 | 0.089196* |
| MAB_4693 | | Putative cytochrome p450 | -5.049168088 | 5.233188127 | 0.18402* |
| MAB_4108c | *rmt4/ elmMIII* | Macrocin-O-methyltransferase MtfB | -5.04486009 | 5.057933 | 0.01373* |

* Not statistically significant
**Not statistically significant based on p-adj value

**Table S5. Strains and plasmids used in this study.**

| Strain/Plasmid | Relevant characteristics | Source |
|---|---|---|
| *Strains* | | |
| *E. coli* | | |
| DH5α | supE44 DlacU169 (f80 lacZDM15) hsdR17recA1 endA1 gyrA69 thi-1relA1 | New England Biolabs |
| *M. abscessus* | | |
| ATCC 19977 | WT parental strain | Kenneth Malcolm, PhD |
| 390S | WT parental strain | Kyle Rohde, PhD |
| 390S Δ*whiB7* | In-frame deletion of *whiB7* | Kyle Rohde, PhD (20) |
| *P. aeruginosa* | | |
| PA14 | WT parental strain | Peter Jorth, PhD |
| *Plasmids* | | |
| pSD5.Hsp60 | Plasmid pSD5 with constitutively expressed Hsp60 promoter | William DePas, PhD |
| pSD5.Hsp60.WhiB7 | Plasmid pSD5.Hsp60 with the *whiB7* coding sequence (+1) | This study |

**Table S6. Primers for cloning and PrimeTime probes for qRT-PCR used in this study.**

**Primers**

| ID | Name | Sequence (5’ to 3’) |
|---|---|---|
| 478117354 | pSD5hsp60_WhiB7_PstI_Front_gib | ATCACTTCCATATGCACTGCAGATGATGACCGTTGAAGTGGAGGCC |
| 478117355 | pSD5hsp60_WhiB7_MluI_Back_gib | TCCATTGAAGACCGGGCCAGAACGCGTTCATGCCGCGGCGGTG |
| 478442339 | pSD5hsp60_seq_F | ACCCGGTGACCTAGACACAT |
| 478442338 | pSD5hsp60_seq_R | CCGACAATGACAACAACCAT |

**PrimeTime Probe and Primer sets**

| Gene target | ID | Name | Sequence (5’ to 3’) |
|---|---|---|---|
| *rpoB* | 473018349 | Mabsc.RpoB set 5 Primer 1 | GACGAGTGCAAAGACAAGGA |
| | 473018350 | Mabsc.RpoB set 5 Primer 2 | CGGGAAATCACCCATGAAGA |
| | 473018351 | Mabsc.RpoB set 5 Probe 5' 6-FAM/ZEN/3' IBFQ | /56-FAM/TCACGGCCG/ZEN/AGTTCATCAACAACA/3IABkFQ/ |
| *whiB7* | 473018345 | WhiB7 set 1 Primer 1 | CTGTGGTTCGCGGAAAG |
| | 473018346 | WhiB7 set 1 Primer 2 | CCCTGCTCAAGAATCTCAC |
| | 473018347 | WhiB7 set 1Probe 5' 6-FAM/ZEN/3' IBFQ | /56-FAM/CCAGGCACT/ZEN/GCGACCGGATC/3IABkFQ/ |

**Plasmid construct**

| Construct | Primer Pair | Vector |
|---|---|---|
| pSD5.Hsp60.WhiB7 | 478117354 - 478117355 | pSD5.Hsp60 |
